# Visual context modulates action perception in 10-month-old infants

**DOI:** 10.1101/131524

**Authors:** Cathleen Bache, Hannes Noack, Anne Springer, Waltraud Stadler, Franziska Kopp, Ulman Lindenberger, Markus Werkle-Bergner

## Abstract

Research on early action perception has documented infants’ astounding abilities in tracking, predicting, and understanding other people’s actions. Common interpretations of previous findings tend to generalize across a wide range of action stimuli and contexts. In this study, ten-month-old infants repeatedly watched a video of a same-aged crawling baby that was transiently occluded. The video was presented in alternation with videos displaying visually either dissimilar movements (i.e., distorted human, continuous object, and distorted object movements) or similar movements (i.e., delayed or forwarded versions of the crawling video). Eye-tracking behavior and rhythmic neural activity, reflecting attention (posterior alpha), memory (frontal theta), and sensorimotor simulation (central alpha), were concurrently assessed. Results indicate that, when the very same movement was presented in a dissimilar context, it was tracked at more rear parts of the target and posterior alpha activity was elevated, suggesting higher demands on attention-controlled information processing. We conclude that early action perception is not immutable but shaped by the immediate visual context in which it appears, presumably reflecting infants’ ability to flexibly adjust stimulus processing to situational affordances.

## Introduction

At the end of the first year of life, infants display remarkable abilities in *action perception*: They pursue and predict another person’s action even if the action is transiently occluded from sight (Bache et al., submitted; Green, Kochukhova, & Gredebäck, 2014; Stapel, Hunnius, Meyer, & Bekkering, 2016). Research has shown that adults’ action perception is not immutable but sensitive to task instruction (e.g., Ocampo & Kritikos, 2010) and immediate visual context (e.g., Güldenpenning, Braun, Machlitt, & Schack, 2015). However, the effects of task, stimulus, and context properties on infants’ action perception have received only little attention (cf. Daum, Gampe, Wronski, & Attig, in press).

Though infants cannot be instructed to watch actions one or another way, action stimuli presented alternately within an experimental session may provide for a visual context that shapes their processing. In line with this view, infants’ action prediction deteriorated when multiple grasping actions were presented successively in contrast to repeating only one grasping action (Henrichs, Elsner, Elsner, Wilkinson, & Gredebäck, 2014). It remains unclear whether, for example, the processing of a crawling movement presented alternately with walking (Stapel, Hunnius, van Elk, & Bekkering, 2010) resembles that presented alternately with object and distorted movements (Bache et al., 2015). However, such modifiable action perception would challenge the integrative interpretation of previous findings.

The present study addresses this issue reanalyzing data from two experiments previously conducted in our lab. In both experiments, 10-month-old infants, who were able to crawl, watched an identical *target video* showing a same-aged crawling baby (i.e., continuous human movement) that was transiently occluded. In Experiment 1, target videos were alternated with videos showing distorted crawling as well as continuous and distorted object movements that were assumed to provide a visually *dissimilar* context (see Bache et al., 2015). In Experiment 2, target videos were alternated with videos showing the same crawling movement in a temporally shifted manner, considered to provide a visually *similar* context (see Bache et al., submitted). Here, we present eye-tracking (ET) and electroencephalographic (EEG) data that indicate differential processing of the very same action when embedded in different visual contexts.

Transient occlusion allows studying processes related to both the external representation of the action during visual perception and the internal representation of the action during occlusion. We were specifically interested in context effects on *attentional*, *mnemonic*, and *sensorimotor* processes, which have been shown to be recruited during the perception of visible and occluded actions (e.g., Bache et al., 2015; Bache et al., submitted; Rotem-Kohavi et al., 2014; Warreyn et al., 2013).

To assess context effects, tracking behavior (via ET) and rhythmic neural activity (via EEG) in theta (4–6 Hz) and alpha (6–9 Hz) frequency ranges were measured simultaneously. The gaze position relative to the target position was taken to reflect tracking accuracy (see Bache et al., submitted for further details). Modulations of frontal theta activity were taken to index mnemonic functions. Sensorimotor simulation was assumed to modulate central alpha activity, while attentional engagement should affect posterior alpha rhythms (see Bache et al., 2015 for further details). As the target action stimuli (i.e., videos) were identical in both experiments, differences in tracking and neural patterns cannot be attributed to perceptual differences but rather to the visual context provided within the experiments.

In a dissimilar context, tracking the target video is likely more demanding and therefore less accurate and proactive (i.e., at more rear parts on the target; Henrichs et al., 2014). Similarly, increased attentional demands (i.e., modulation of posterior alpha activity) are expected (Klimesch, 2012). By contrast, similar contexts permit more fine-grained action prediction processes (i.e., modulations of central alpha activity; Wolpert & Flanagan, 2001).

## Methods

### Participants

Two experiments were conducted at the Max Planck Institute for Human Development, Berlin, Germany. Seventy-nine and 99 10-month-old infants (± 10 days) participated in Experiment 1 and Experiment 2, respectively. For the purpose of this study, participants were randomly assigned to one of *three experimental groups* watching a target video (i.e., crawling) alternately with either dissimilar (Experiment 1: *Context Dissimilar*) or similar movements (Experiment 2: *Context Similar Delay* vs. *Context Similar Forward*). The experiments were approved by the Institute’s Ethics Committee.

Participants were recruited via the Institute’s database, consisting of parents interested in participating in infant studies, according to the following criteria: the infant *(a)* was born at term (week of gestation ≥ 37, birth weight ≥ 2500 g), and according to the parents *(b)* had no visual impairments or health issues, and *(c)* was capable of crawling on hands and knees with the stomach lifted while not yet walking. Participants were excluded from ET, EEG, or both analyses for the following reasons: *(a)* following preparation of EEG and ET, the infant was too fussy to be tested (n = 5 in Experiment 1/n = 8 in Experiment 2), *(b)* the infant did not crawl a distance of 1.5 m in the lab (n = 4/4), *(c)* there were technical issues with video, ET, and/or EEG recording (n = 8/17), *(d)* the calibration failed (n = 1/3), and the infant provided *(e)* less than 10 % of artifact-free *ET trials* of the actually watched trials (n = 5/1), and *(g)* less than 10 artifact-free *EEG trials* per condition (n = 29/37).

The final *ET sample* consisted of *(a)* **42** infants in Context Dissimilar, *(b)* **32** infants in Context Similar Delay, and *(c)* **31** infants in Context Similar Forward. The final *EEG sample* comprised *(a)* **25** infants in Context Dissimilar, *(b)* **24** infants in Context Similar Delay, and *(c)* **25** infants in Context Similar Forward. Descriptive information on the ET and EEG samples are provided in Table 1.

**Table 1.**
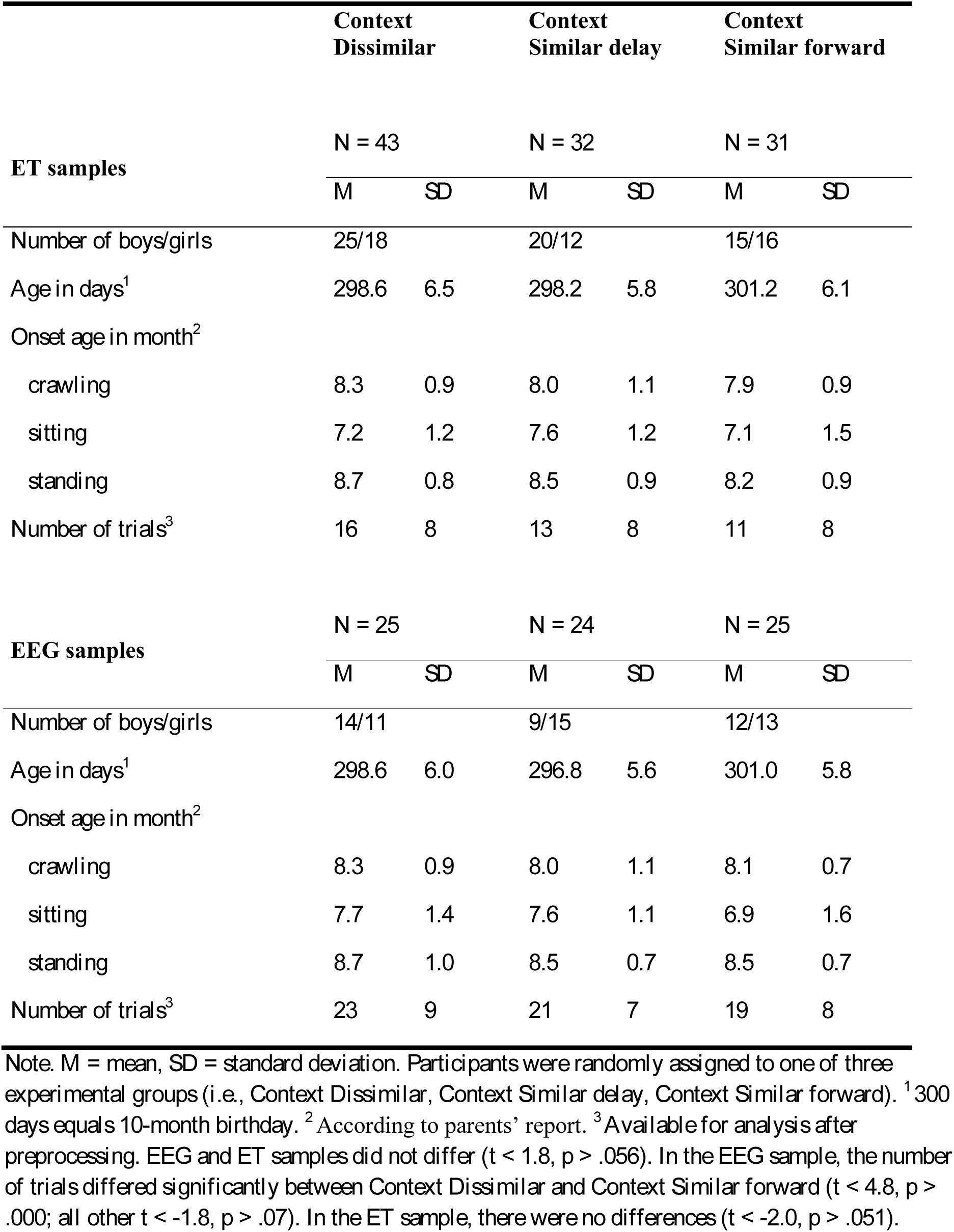
Descriptive information on the ET and EEG samples in all experimental groups

|  | **Context**<br>**Dissimilar** |  | **Context**<br>**Similar delay** |  | **Context**<br>**Similar forward** |  |
| --- | --- | --- | --- | --- | --- | --- |
|  | N = 43 |  | N = 32 |  | N = 31 |  |
| **ET samples** | *M* | *SD* | *M* | *SD* | *M* | *SD* |
| Number of boys/girls | 25/18 |  | 20/12 |  | 15/16 |  |
| Age in days<sup>1</sup> | 298.6 | 6.5 | 298.2 | 5.8 | 301.2 | 6.1 |
| Onset age in month<sup>2</sup> |  |  |  |  |  |  |
| crawling | 8.3 | 0.9 | 8.0 | 1.1 | 7.9 | 0.9 |
| sitting | 7.2 | 1.2 | 7.6 | 1.2 | 7.1 | 1.5 |
| standing | 8.7 | 0.8 | 8.5 | 0.9 | 8.2 | 0.9 |
| Number of trials<sup>3</sup> | 16 | 8 | 13 | 8 | 11 | 8 |
| **EEG samples** | N = 25 |  | N = 24 |  | N = 25 |  |
|  | *M* | *SD* | *M* | *SD* | *M* | *SD* |
| Number of boys/girls | 14/11 |  | 9/15 |  | 12/13 |  |
| Age in days<sup>1</sup> | 298.6 | 6.0 | 296.8 | 5.6 | 301.0 | 5.8 |
| Onset age in month<sup>2</sup> |  |  |  |  |  |  |
| crawling | 8.3 | 0.9 | 8.0 | 1.1 | 8.1 | 0.7 |
| sitting | 7.7 | 1.4 | 7.6 | 1.1 | 6.9 | 1.6 |
| standing | 8.7 | 1.0 | 8.5 | 0.7 | 8.5 | 0.7 |
| Number of trials<sup>3</sup> | 23 | 9 | 21 | 7 | 19 | 8 |

*Note.* M = mean, SD = standard deviation. Participants were randomly assigned to one of three experimental groups (i.e., Context Dissimilar, Context Similar delay, Context Similar forward). <sup>1</sup> 300 days equals 10-month birthday. <sup>2</sup> According to parents' report. <sup>3</sup> Available for analysis after preprocessing. EEG and ET samples did not differ ( $t < 1.8$ , $p > .056$ ). In the EEG sample, the number of trials differed significantly between Context Dissimilar and Context Similar forward ( $t < 4.8$ , $p > .000$ ; all other $t < -1.8$ , $p > .07$ ). In the ET sample, there were no differences ( $t < -2.0$ , $p > .051$ ).

Not all infants contributed data to both measures, and artifact-free trials were randomly distributed across the experiment. High attrition is commonly observed in studies simultaneously measuring both brain and eye data (e.g., Haith, 2004; Stets, Stahl, & Reid, 2012). Therefore, separate analyses were performed for ET and EEG data (Bache et al., submitted; Stapel et al., 2010). As a further consequence, a systematic analysis of data over experimental time was not conducted, because it would have required reducing the number of available trials and participants substantially.

### Stimulus material and procedure

Participants repeatedly watched the *target video* of a same-aged baby continuously crawling in front of a gray background, which was transiently occluded. The presentation of the target video alternated with further videos assumed to constitute a certain visual context to the target video.

In Experiment 1, participants additionally saw videos displaying distorted crawling movement, continuous object movement, and distorted object movement in a within-subjects design (Bache et al., 2015)^1^. Thus, the videos varied both the stimulus target (i.e. baby vs. object) and the smoothness of the movement (i.e., continuous vs. distorted), taken to constitute a visually dissimilar context (*Context Dissimilar*).

In Experiment 2, participants additionally watched videos of the same movement (i.e., crawling baby) differing subtly in the timing following the occlusion (Bache et al., submitted). That is the timing was continuous in the target video (i.e. movement continued correctly in time during occlusion) and, in a between-subjects design, either delayed (*Context Similar Delay*) or forwarded (*Context Similar Forward*) in the other videos. Accordingly, the visual context of the target video was considered highly similar.

In each video, the ongoing movement (2480 ms of pre-occlusion) was transiently occluded by a full-screen black occlusion (520 ms of occlusion) and continued subsequently (1000 ms of post-occlusion). Movements were presented from both left to right and right to left (i.e., flipped versions of the original videos were shown). Conditions were presented in blocks of six repetitions; the order of blocks was quasi-randomized. A maximum of 24 blocks were presented depending on the infants’ compliance. More detailed information on stimuli and procedure are provided in Bache et al. (2015; submitted). In sum, the present report covers data collected in two experiments comparing three experimental groups of 10-month-olds watching a target video in different visual contexts.

### Data recording and analysis

ET data were recorded continuously at 250 Hz using an EyeLink 1000 remote system eye-tracker (SR research, Ottawa, Canada). Simultaneously, EEG was recorded continuously from 32 active electrodes (actiCap by BrainProducts) at 1000 Hz using a BrainAmp DC amplifier (BrainProducts GmbH, Gilching, Germany; further specifications: online pass-band 0.1–250 Hz, right mastoid reference, and ground at AFz, offline linked-mastoids reference).

*ET data* were visually inspected for measurement error and compliance failure to determine clean trials for analyses. We focused on gaze positions on the x-dimension (Gx) as the crawling movement progressed on the horizontal dimension. For each measurement point, the distance between gaze position and mean target position (i.e., center of mass) was calculated to relate tracking behavior to target movement over time. For statistical analysis, the *mean distance* was calculated across 500 ms time windows separately for each phase of the target video (i.e., last 500 ms of pre-occlusion vs. 500 ms of occlusion vs. first 500 ms of post-occlusion). More detailed information on the preprocessing and analysis of ET data is provided in Bache et al. (submitted).

*EEG data* were visually inspected and cleaned from artifacts using Independent Component Analysis (ICA; Jung et al., 2000). Rhythmic neural activity was analyzed by means of fast Fourier transformation (FFT) applying an ideographic filter approach (e.g., Grandy et al., 2013; see also Karch, Sander, von Oertzen, Brandmaier, & Werkle-Bergner, 2015; Nesselroade, Gerstorf, Hardy, & Ram, 2007). In line with the literature, *frontal theta* activity was defined as oscillatory activity within 4-6 Hz at frontal electrodes (Orekhova, Stroganova, & Posikera, 1999). *Alpha* activity was defined as oscillatory activity within 6-9 Hz at *central* and *posterior* electrodes (Marshall, Bar-Haim, & Fox, 2002; Marshall, Young, & Meltzoff, 2011). More detailed information on EEG preprocessing and analysis is provided in Bache et al. (2015).

Only data from target videos were statistically compared. Analyses were performed in SPSS (SPSS Inc., 1989-2006, USA) using mixed effects repeated-measures ANOVAs with between-subject factor Context (Context Dissimilar vs. Context Similar Delay vs. Context Similar Forward) and within-subjects factor Phase (pre-occlusion vs. occlusion vs. post-occlusion) separately for each measure of eye movement (i.e., *mean distance*) and rhythmic neural activity (i.e., *frontal theta*, *central alpha*, and *posterior alpha*).

## Results

### Eye-tracking data

***Mean distance per phase***

*Figure 1* depicts the mean horizontal distance between gaze position and mean target position over time. A mixed ANOVA revealed significant main effects of *(a)* Phase (*F*_(1.4, 142.5)_ = 124.56, *p* =.000, 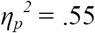)^2^, and *(b)* Context (*F* = 7.25, *p* = .001, *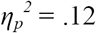*), as well as *(c)* a Phase by Context interaction (*F*_(2.0, 106.5)_ = 4.76, *p* = .001, *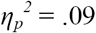*)^3^. An overview of the results is provided in *Figure 2*.

**Figure 1.**
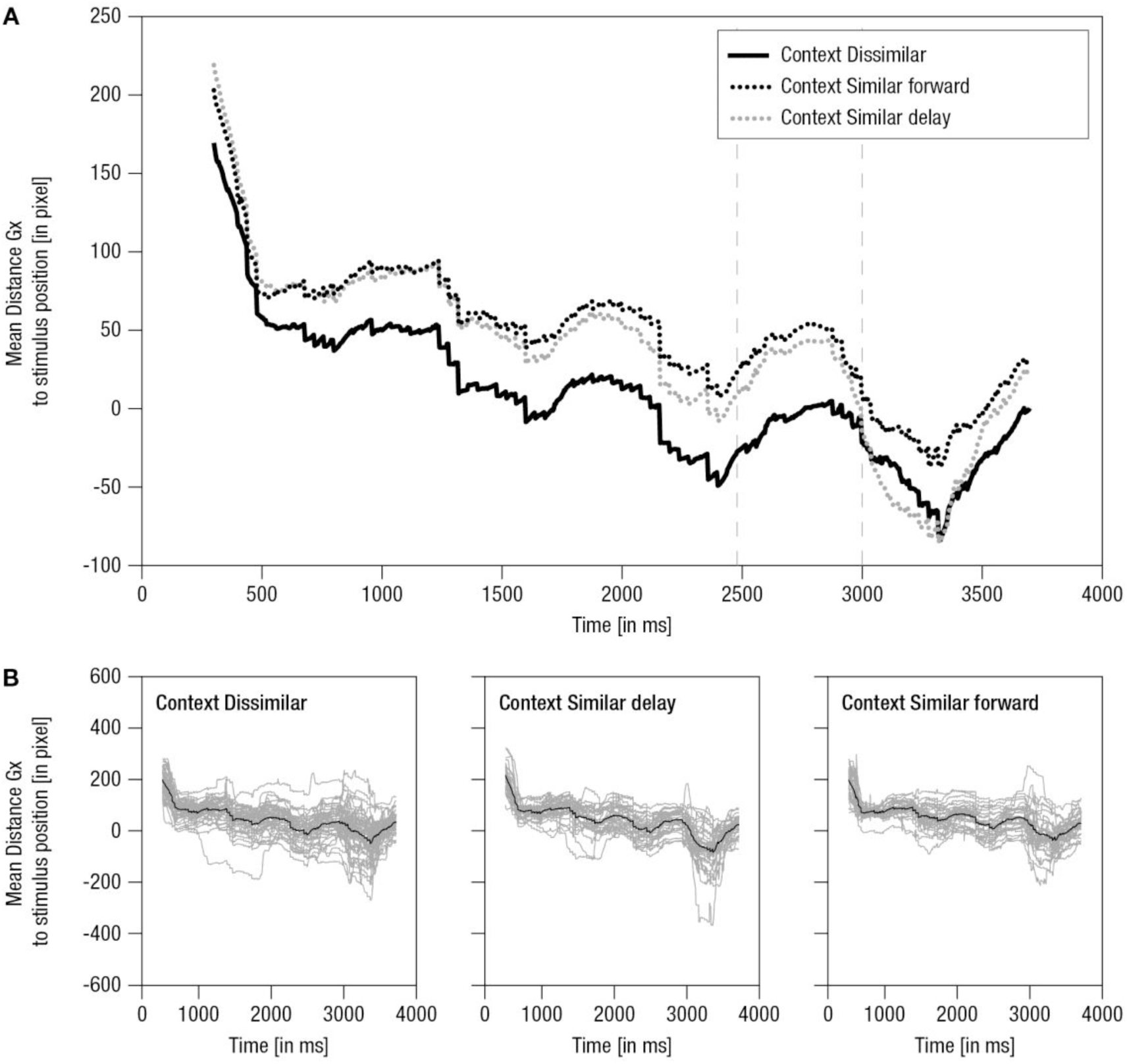
Mean horizontal distance between gaze position and mean stimulus position over time. A) Grand averaged distance over time. Lines: Solid black – Context Dissimilar, Pointed black – Context Similar delay, Pointed gray – Context Similar forward, Vertical – occlusion on- and off-set. B) Single averaged distance over time (gray) for the final sample for ET analysis including respective grand average (black): Left – Context Dissimilar, Middle – Context Similar delay, Right – Context Similar forward. Note, average stimulus dimension: 281 pixel.

**Figure 2.**
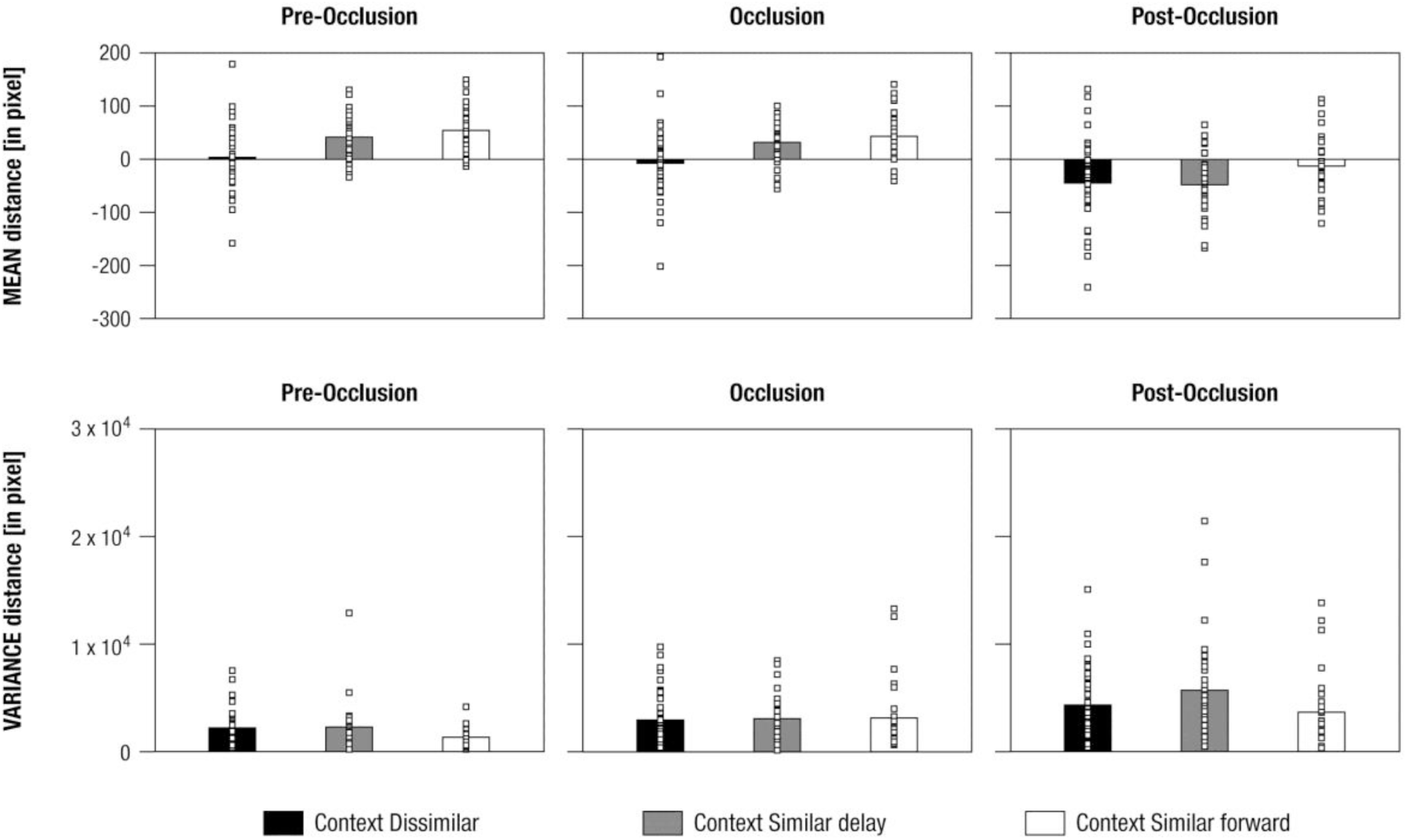
Mean differences in distance in response to human continuous movement across visual contexts (i.e., Context Dissimilar, Context Similar delay, Context Similar forward) and phases (i.e., pre-occlusion, occlusion, and post-occlusion) for mean distance between gaze positions and mean stimulus positions. Squares indicate single cases demonstrating the distribution within the sample.

To follow-up on the interaction effect, independent *t*-Tests per level of Phase were calculated showing that:

*(1) during pre-occlusion and occlusion*, the target was tracked approximately 40 pixel more backwards (i.e., gaze on mid to rear parts of target, such as bottom or knees) in *Context Dissimilar* (pre-occlusion: *M* = 3.96, *SE* = 9.13, occlusion: *M* = -7.63, *SE* = 9.85) than in *Context Similar Delay* (pre-occlusion: *M* = 41.56, *SE* = 7.7, occlusion: *M* = 31.58, *SE* = 9.85; *t*_(72)_ > 3.03, *p* < .003) as well as in *Context Similar Forward* (pre-occlusion: *M* = 54.22, *SE* = 7.84, occlusion: *M* = 42.97, *SE* = 8.4; *t*_(71)_ > 3.7, *p* < .000), whereas Context Similar Delay and Context Similar Forward did not differ (*t*_(61)_ < -1.05, *p* > .257);

*(2) during post-occlusion*, though the target was tracked approximately 30 pixel more forwards (i.e. gaze on mid parts) in *Context Similar Forward* (*M* = -12.91, *SE* = 10.34) than in *Context Dissimilar* and *Context Similar Delay*, the differences did not reach the Bonferroni-corrected significance level (*t* < 1.93, *p* > .048). *Context Dissimilar* (*M* = -44.53, *SE* = 11.81) and *Context Similar Delay* (*M* = -47.88, *SE* = 9.9) did not differ in mean distance (*t*_(72)_ = -0.28, *p* = .836).

Though the mean distance differed, tracking patterns were highly systematic across contexts during the pre-occlusion and occlusion phases (see *Figure 1*). To further evaluate the differences in mean distance at the *transition* from occlusion to post-occlusion, the absolute difference of mean distance in occlusion and post-occlusion time windows was calculated for each context. Independent samples *t*-Tests showed that in *Context Similar Delay* (*M* = 80.16, *SE* = 9.25), compared to the actual stimulus movement, tracking slowed down more (i.e., was more reactive) than in Context Dissimilar (*M* = 52.19, *SE* = 5.50; *t*_(72)_ = 2.73, *p* = .008). The difference between Context Similar Delay and Context Similar Forward (*M* = 57.54, *SE* = 6.6; *t*_(61)_ = 1.98, *p* = .025) did not reach Bonferroni-corrected significance. ‘Slowing down’ did not differ between Context Similar Forward and Context Dissimilar (*t*_(71)_ = 0.62, *p* = .533).^4^

In sum, infants’ tracking behavior during pre-occlusion and occlusion was more conservative (i.e., at more mid to rear parts) when the target video was presented in alternation with dissimilar movements. This supports the hypothesis that the tracking of an action is less accurate and less proactive in visually dissimilar contexts. Furthermore, following reappearance, infants rather reactively followed the target (i.e., slowing down of tracking), when it was alternated with delayed versions of movement (i.e., similar context) than with dissimilar movements. Hence, even in highly similar contexts, action perception appeared context-dependent. Together, results suggest that tracking the very same human movement depended on the visual context provided within the experimental session.

### EEG data

#### Posterior alpha activity

A mixed ANOVA for posterior alpha activity showed a significant main effect of Phase (*F*_(1.7, 35.22)_ = 92.40, *p* = .000, *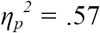*) and a significant interaction effect of Phase and Context (*F*_(4, 4.1)_ = 5.33, *p* = .000, 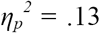). The main effect of Context did not reach significance (*F*_(2, 4.84)_ = 2.2, *p* = .117, *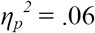*). *Figure 3* provides an overview of the FFT results.

**Figure 3.**
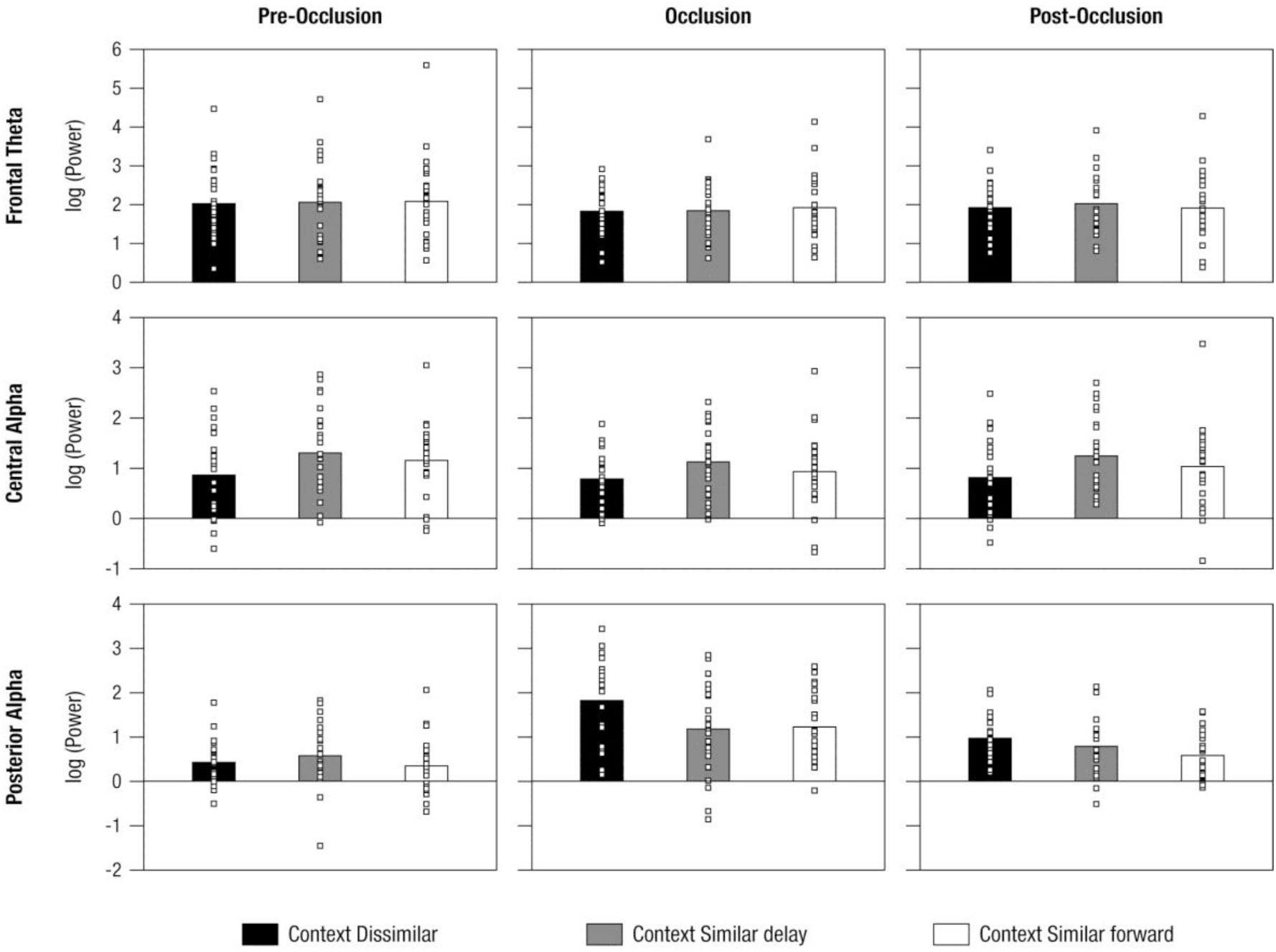
Mean power differences in response to human continuous movement across visual contexts (i.e., Context Dissimilar, Context Similar delay, Context Similar forward) and phases (i.e., pre-occlusion, occlusion, and post-occlusion) for frontal theta, central alpha, and posterior alpha activity. Squares indicate single cases demonstrating the distribution within the sample.

Following up on the interaction effect, applying univariate ANOVAs separately per level of Phase, a main effect of Context was found for *occlusion* (*F*_(2, 6.39)_ = 3.91, *p* = .025, *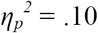*) and *post-occlusion* (*F*_(2, 1.86)_ = 3.24, *p* = .045, *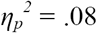*) but not for pre-occlusion (*F* = 0.89, *p* = .416, *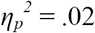*) phase. Separate independent-samples *t*-Tests revealed that:

*(1) during occlusion*, posterior alpha activity was higher in *Context Dissimilar* (*M* = 1.82, *SE* = .18) than in *Context Similar Delay* (*M* = 1.18, *SE* = .21; *t* = 2.30, *p* = .026) as well as in *Context Similar Forward* (*M* = 1.23, *SE* = .15; *t* = 2.54, *p* = .014), whereas similar contexts did not differ from each other (*t* = -1.18 and *p* = .86);

(2) *during post-occlusion*, posterior alpha activity was significantly higher in *Context Dissimilar* (*M* = 0.97, *SE* = .09) than *Context Similar Forward* (*M* = 0.57, *SE* = .09; *t* = 2.80, *p* = .007; other *t* ≤ 1.292 and *p* ≥ .203).

To better understand the timing of posterior alpha modulations during occlusion and post-occlusion, additional analyses of time-course effects were performed. To this end, the pre-processed EEG data were band-pass filtered at the individual peak frequency (± 1 Hz), and amplitude time-courses were extracted by taking the absolute values of the Hilbert-transformed time-series (Le Van Quyen et al., 2001). In doing so, leakage in the frequency domain was attenuated, though smearing in the time domain cannot be prevented (Widmann, Schroger, & Maess, 2015). Amplitude values at each measurement point were statistically compared using independent *t*-Tests. To use a conservative threshold, results were only considered reliable when at least 10 consecutive measurement points (i.e., 100 ms) met an an alpha level of .01. The results suggest that only *Context Dissimilar* and *Context Similar Delay* differed significantly from one another, and that effects can be attributed to the occlusion rather than to the post-occlusion phase (see *Figure 4*). In sum, neural activity related to attentional demands was increased during the occlusion of an ongoing crawling movement when infants observed this crawling in the context of dissimilar trials.

**Figure 4.**
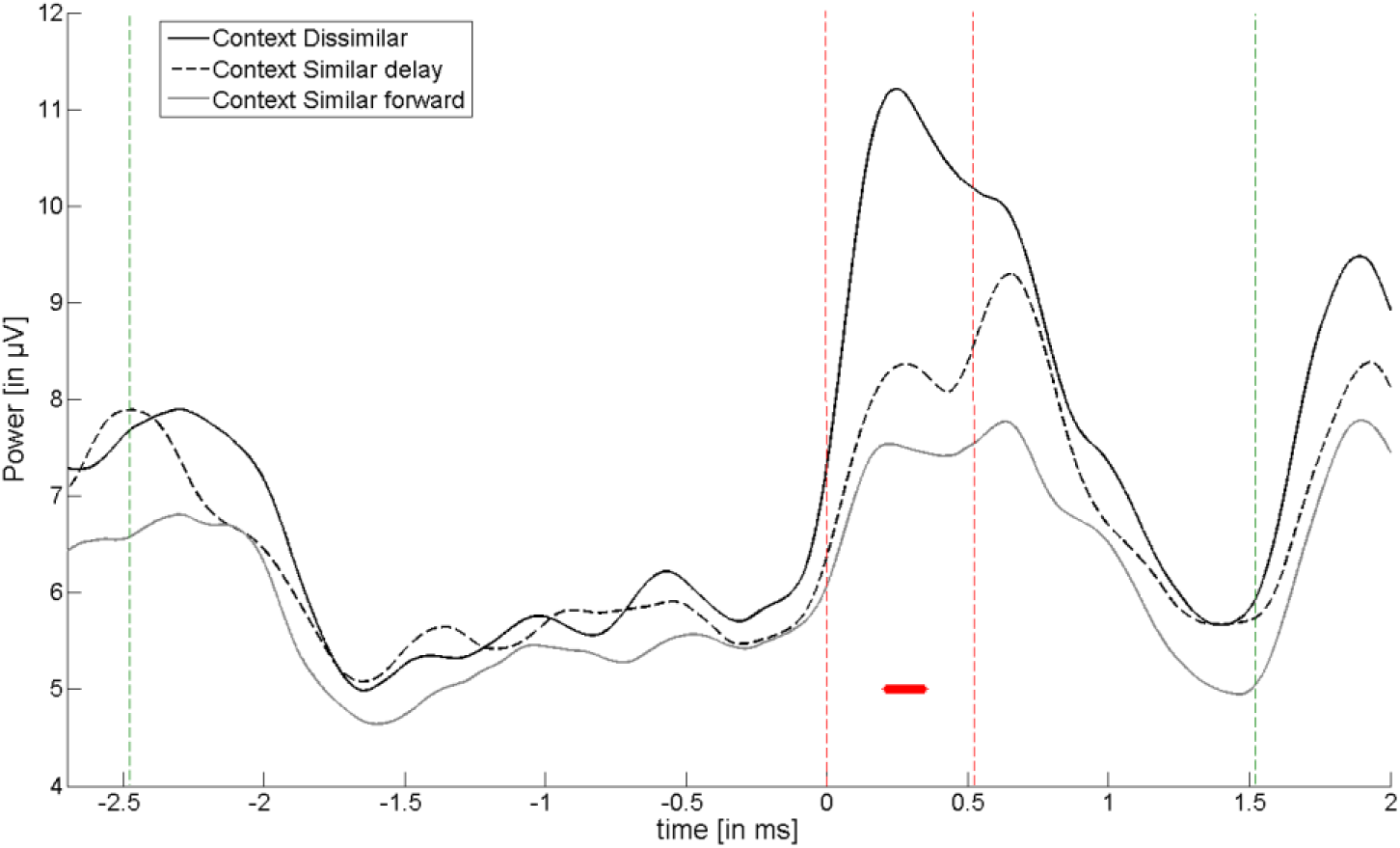
Mean power differences in posterior alpha over time in response to human continuous movement across visual contexts. Lines: Solid black – Context Dissimilar, Dashed black – Context Similar delay, Gray – Context Similar forward, Vertical red – on- and offset of occlusion, Vertical green – trial on- and offset, Red crosses indicate significant differences in *t*-values (plotted if *p* < .01 in at least 10 consecutive time points).

#### Frontal theta and central alpha activity

Using mixed ANOVA, there were main effects of Phase for frontal theta (*F*_(1.6, 1.4)_ = 6.49, *p* =.004, *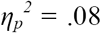*) and central alpha activity (*F*_(1.6, 1.4)_ = 6.49, *p* = .004, *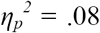*) without evidence for further effects (frontal theta: all *F* < 0.56; all *p >* .692; central alpha: (*F* < 2.13; *p >* .13). Thus, frontal theta activity, representing processes related to working memory, did not differ between the visual contexts. Similarly, central alpha activity, serving as indicator of sensorimotor processing, did not differentiate between similar and dissimilar contexts.

## Discussion

This study investigated effects of the visual context on attentional, mnemonic, and sensorimotor processing during action perception. In three experimental groups, 10-month-old infants observed the very same transiently occluded crawling movement. We examined whether tracking behavior and associated neural activity in response to this target video were influenced by the alternated presentation of visually either similar or dissimilar movements. The results showed that infants tracked the target video less accurately and proactively in a dissimilar than in a similar context, presumably reflecting increased attentional demands under low-similarity conditions. Moreover, tracking was less accurate following reappearance, when the target video was alternated with similar but temporally delayed versions of the very movement. In sum, our results show that infants’ action perception processes were modified by visual context.

To delineate these context effects, posterior alpha activity was assessed as a marker of attention (Klimesch, 2012). Results showed that the strongest alpha activity occurred with a dissimilar context during occlusion. This is in line with the notion of *active inhibition* during controlled information processing in visual areas of the brain when internal cognitive operations, such as maintaining a target during occlusion, are executed at non-perceptual areas in the brain (e.g., Jensen & Mazaheri, 2010). Hence, a dissimilar context probably increased demands on controlling target maintenance resulting in increased inhibition of potentially interfering information. Alternatively, modulations in posterior alpha activity may reflect the use of visual *imagery* during occlusion in which primary visual areas are also involved (Kosslyn, Ganis, & Thompson, 2001). From this perspective, embedding the target video in similar contexts may have facilitated actively imagining the occluded action resulting in relatively reduced alpha activity (Hanslmayr, Staudigl, & Fellner, 2012; Werkle-Bergner et al., 2014).

Corresponding to neural modulations, infants tracked the target at more rear parts, and thus more reactively, prior to its disappearance when it was presented in a dissimilar context. Together, the findings therefore suggest that attentional demands on perceptual processing increased and shaped tracking behavior when the target video was alternated with visually dissimilar movements.

The present context effects may emerge when a *prime* perceived prior to the target video conveys corresponding information and thereby facilitates the processing of the target (Posner, 1994). Motion is a very potent prime in adults (e.g., Güldenpenning et al., 2015) and infants (Gredebäck & Daum, 2015). Accordingly, similar contextual trials may have constituted a prime to the target video which allowed infants to track the target at more front parts prior to occlusion. As the prime was not presented immediately preceding the target video, the present findings suggest that infants take distant information into account – the context of an entire experimental session – when processing a transiently occluded crawling movement.

Action perception may however not only be modified by obvious visual differences in stimulus presentation, but also by rather subtle task requirements. In the dissimilar context, the target video could be differentiated on the basis of perceptually analyzing target (i.e., baby vs. object) and movement (continuous vs. distorted) properties (Bache et al., 2015). In contrast, in the similar contexts, the target video could only be differentiated on the basis of timed internal representations during occlusion (Bache et al., submitted; Graf et al., 2007). Importantly, presented in alternation with delayed videos, the tracking of the target video slowed down considerably following occlusion, suggesting that the presumably timed internal representation was too fast. This finding corresponds to evidence suggesting that delayed and forwarded time-shifts may not be processed similarly (Bache et al., submitted), possibly because infants learned to accommodate to delayed timing during everyday mother-infant interaction (Striano, Henning, & Stahl, 2006).

Unfortunately, the high prevalence of missing EEG and ET data does not allow us to analyze the time course of context effects (cf. Stets & Reid, 2011). Furthermore, because we did not present the target video without visual context, we are unable to determine the contextual processing in relation to baseline action perception. Nevertheless, and irrespective of whether contextual information facilitates or misleads processing, the present results suggest that early action perception is flexible and allows for adapting to changing situational constraints (Schoner, Dijkstra, & Jeka, 1998; Skerry, Carey, & Spelke, 2013).

In addition to attention, processes related to *sensorimotor simulation* (i.e., central alpha activity) and *memory* (i.e., frontal theta activity) were investigated. Central alpha activity did not differ between visual contexts. Hence, we did not find evidence that sensorimotor processing, serving predictive functions (Wolpert & Flanagan, 2001), differentially facilitated action perception in either of the visual contexts investigated here. Furthermore, frontal theta activity did not differentiate between visual contexts. As working memory content is permanently updated, this finding may indicate that maintaining and integrating extracted information across time and space was equally demanding in the visual contexts contrasted here (Simons & Spiers, 2003). The contextual effects observed here may be specific to transiently occluded actions as previous research showed that overall attentional demands increase in occlusion events (e.g., Moore, Borton, & Darby, 1978).

In sum, we conclude that the perception of a crawling movement is not immutable but influenced by visual context. Even though infants were not prompted to watch the identical target video in one or another way, the visual context promoted differential processing. Hence, generalization based on a specific set of action stimuli used in a given study requires caution, as the same set of stimuli may have evoke a different response if the context was different (Poulton, 1973). Conversely, the systematic variation of contextual conditions can be seen as an experimental manipulation for characterizing the flexibility of infants’ minds.

## Acknowledgements

This research was supported by the Max Planck Research Network for the Cognitive and Neurosciences (Maxnet *Cognition*) and funded by the Max Planck Society. The study was conducted in partial fulfillment of the doctoral dissertation of CB. CB received training and financial support from the International Max Planck Research School on the Life Course (*LIFE*, <u>http://www.imprs-life.mpg.de/en</u>). WS received funding of Deutsche Forschungsgemeinschaft, DFG (Project: STA 1076/1-1). MW-B’s work is supported by a grant from the German Research Foundation (DFG, WE 4269/3-1) as well as an *Early Career Research Fellowship 2017 – 2019* awarded by the Jacobs Foundation. We cordially thank the infants and their parents for participating in this study and our student assistants for their support in data collection, coding and preprocessing. We further wish to thank Berndt Wischnewski for technical support.

## Footnotes

1 Note, in Experiment 1, the test session was repeated after 2-10 days to obtain sufficient artifact-free trials in all conditions.

2 Greenhouse-Geisser corrections were applied whenever indicated.

3 We also analyzed *variance in distance* to determine tracking consistency across participants (see Bache et al., submitted). There were no significant differences in variance across visual contexts.

4 The number of infants differed between experimental groups with higher sample size in Context Dissimilar compared to both Context Similar Delay and Context Similar Forward (see *Participants*). As significant differences did not yield a similar pattern across phases, differences in sample size do probably not account for the found effects.

## References

Bache, C., Kopp, F., Springer, A., Stadler, W., Lindenberger, U., & Werkle-Bergner, M. (2015). Rhythmic neural activity indicates the contribution of attention and memory to the processing of occluded movements in 10-month-old infants. International Journal of Psychophysiology, 98, 201–212.

Bache, C., Springer, A., Noack, H., Stadler, W., Kopp, F., Lindenberger, U., et al. (submitted). 10-month-old infants are sensitive to the time course of perceived actions: evidence from a study combining eye-tracking and EEG. Frontiers in Human Neuroscience.

Daum, M. M., Gampe, A., Wronski, C., & Attig, M. (in press). Effects of movement distance, duration, velocity, and type on action prediction in 12-month-olds. Infant Behavior & Development.

Graf, M., Reitzner, B., Corves, C., Casile, A., Giese, M., & Prinz, W. (2007). Predicting point-light actions in real-time. NeuroImage, 36, 22–32.

Grandy, T. H., Werkle-Bergner, M., Chicherio, C., Schmiedek, F., Lovden, M., & Lindenberger, U. (2013). Peak individual alpha frequency qualifies as a stable neurophysiological trait marker in healthy younger and older adults. Psychophysiology, 50(6), 570–582.

Gredebäck, G., & Daum, M. M. (2015). The Microstructure of Action Perception in Infancy: Decomposing the Temporal Structure of Social Information Processing. Child Development Perspectives, 9(2), 79–83.

Green, D., Kochukhova, O., & Gredebäck, G. (2014). Extrapolation and direct matching mediate anticipation in infancy. Infant Behavior & Development, 37(1), 111–118.

Güldenpenning, I., Braun, J. F., Machlitt, D., & Schack, T. (2015). Masked priming of complex movements: perceptual and motor processes in unconscious action perception. Psychological Research, 79(5), 801–812.

Haith, M. M. (2004). Progress and standardization in eye movement work with human infants. Infancy, 6(2), 257–265.

Hanslmayr, S., Staudigl, T., & Fellner, M. C. (2012). Oscillatory power decreases and long-term memory: the information via desynchronization hypothesis. Frontiers in Human Neuroscience, 6, 74.

Henrichs, I., Elsner, C., Elsner, B., Wilkinson, N., & Gredebäck, G. (2014). Goal certainty modulates infants’ goal-directed gaze shifts. Developmental Psychology, 50(100-107..

Jensen, O., & Mazaheri, A. (2010). Shaping functional architecture by oscillatory alpha activity: gating by inhibition. Frontiers in Human Neuroscience, 4.

Jung, T. P., Makeig, S., Humphries, C., Lee, T. W., Mckeown, M. J., Iragui, V., et al. (2000). Removing electroencephalographic artifacts by blind source separation. Psychophysiology, 37(2), 163–178.

Karch, J. D., Sander, M. C., von Oertzen, T., Brandmaier, A. M., & Werkle-Bergner, M. (2015). Using within-subject pattern classification to understand lifespan age differences in oscillatory mechanisms of working memory selection and maintenance. NeuroImage, 118, 538–552.

Klimesch, W. (2012). Alpha-band oscillations, attention, and controlled access to stored information. Trends in Cognitive Sciences, 16(12), 606–617.

Kosslyn, S. M., Ganis, G., & Thompson, W. L. (2001). Neural foundations of imagery. Nature reviews. Neuroscience, 2(9), 635–642.

Le Van Quyen, M., Foucher, J., Lachaux, J.-P., Rodriguez, E., Lutz, A., Martinerie, J., et al. (2001). Comparison of hilbert transform and wavelet methods for the analysis of neuronal synchrony. Journal of Neuroscience Methods, 111, 83–98.

Marshall, P. J., Bar-Haim, Y., & Fox, N. A. (2002). Development of the EEG from 5 months to 4 years of age. Clinical Neurophysiology, 113(8), 1199–1208.

Marshall, P. J., Young, T., & Meltzoff, A. N. (2011). Neural correlates of action observation and execution in 14-month-old infants: an event-related EEG desynchronization study. Developmental Science, 14(3), 474–480.

Moore, M. K., Borton, R., & Darby, B. L. (1978). Visual Tracking in Young Infants - Evidence for Object Identity or Object Permanence. Journal of Experimental Child Psychology, 25(2), 183–198.

Nesselroade, J. R., Gerstorf, D., Hardy, S. A., & Ram, N. (2007). Idiographic filters for psychological constructs. Measurement: Interdisciplinary Research and Perspectives (5), 217–235.

Ocampo, B., & Kritikos, A. (2010). Placing actions in context: motor facilitation following observation of identical and non-identical manual acts. Experimental Brain Research, 201(4), 743–751.

Orekhova, E. V., Stroganova, T. A., & Posikera, I. N. (1999). Theta synchronization during sustained anticipatory attention in infants over the second half of the first year of life. International Journal of Psychophysiology, 32(2), 151–172.

Posner, M. I. (1994). Attention: the mechanisms of consciousness. Proceedings of the National Academy of Sciences of the United States of America, 91(16), 7398–7403.

Poulton, E. C. (1973). Unwanted range effects from using within-subject experimental designs. Psychological Bulletin, 80(2), 113–121.

Rotem-Kohavi, N., Hilderman, C. G. E., Liu, A. P., Makan, N., Wang, J. Z., & Virji-Babul, N. (2014). Network analysis of perception-action coupling in infants. Frontiers in Human Neuroscience, 8.

Schoner, G., Dijkstra, T. M. H., & Jeka, J. J. (1998). Action-Perception Patterns Emerge From Coupling and Adaptation. Ecological Psychology, 10(3-4), 323–346.

Simons, J. S., & Spiers, H. J. (2003). Prefrontal and medial temporal lobe interactions in long-term memory. Nature Reviews: Neuroscience, 4(8), 637–648.

Skerry, A. E., Carey, S. E., & Spelke, E. S. (2013). First-person action experience reveals sensitivity to action efficiency in prereaching infants. Proceedings of the National Academy of Sciences of the United States of America, 110(46), 18728–18733.

Stapel, J. C., Hunnius, S., Meyer, M., & Bekkering, H. (2016). Motor system contribution to action prediction: temporal accuracy depends on motor experience. Cognition, 148, 71–78.

Stapel, J. C., Hunnius, S., van Elk, M., & Bekkering, H. (2010). Motor activation during observation of unusual versus ordinary actions in infancy. Social Neuroscience, 5(5-6), 451–460.

Stets, M., & Reid, V. M. (2011). Infant ERP amplitudes change over the course of an experimental session: Implications for cognitive processes and methodology. Brain and Development, 33(7), 558–568.

Stets, M., Stahl, D., & Reid, V. M. (2012). A meta-analysis investigating factors underlying attrition rates in infant ERP studies. Developmental Neuropsychology, 37(3), 226–252.

Striano, T., Henning, A., & Stahl, D. (2006). Sensitivity to interpersonal timing at 3 and 6 months of age. Interaction Studies, 7(2), 251–271.

Warreyn, P., Ruysschaert, L., Wiersema, J. R., Handl, A., Pattyn, G., & Roeyers, H. (2013). Infants’ mu suppression during the observation of real and mimicked goal-directed actions. Developmental Science, 16(2), 173–185.

Werkle-Bergner, M., Grandy, T. H., Chicherio, C., Schmiedek, F., Lövdén, M., & Lindenberger, U. (2014). Coordinated Within-trial Dynamics of Low-Frequency Neural Rhythms Controls Evidence Accumulation. Journal of Neuroscience, 34(25), 8519–8528.

Widmann, A., Schroger, E., & Maess, B. (2015). Digital filter design for electrophysiological data--a practical approach. Journal of neuroscience methods, 250, 34–46.

Wolpert, D. M., & Flanagan, J. R. (2001). Motor prediction. Current Biology, 11(18), 729–732.

